# DNA data (genome skims and metabarcodes) paired with chemical data demonstrate utility for retrospective analysis of forage linked to fatal poisoning of cattle

**DOI:** 10.1101/2025.01.06.631623

**Authors:** Daniel Cook, Brandon Kocurek, Clint Stonecipher, Kevin D. Welch, Dale R. Gardner, Mark Mammel, Elizabeth Reed, Padmini Ramachandran, David Erickson, Seth Commichaux, Andrea Ottesen

## Abstract

Prepared and stored feeds, fodder, silage, and hay may be contaminated by toxic plants resulting in the loss of livestock. Several poisonous plants have played significant roles in livestock deaths from forage consumption in recent years in the Western United States including *Salvia reflexa*. Metagenomic data, genome skims and metabarcodes, have been used for identification and characterization of plants in complex matrices including diet composition of animals, mixed forages, and herbal products. Here, chemistry, genome skims, and metabarcoding were used to retrospectively describe the composition of contaminated alfalfa hay from a case of *Salvia reflexa* poisoning that killed 165 cattle. Genome skims and metabarcoding provided similar estimates of the relative abundance of the *Salvia* in the hay samples when compared to chemical methods. Additionally, genome skims and metabarcoding provided similar estimates of species composition in the contaminated hay and rumen contents of poisoned animals. The data demonstrate that genome skims and DNA metabarcoding may provide useful tools for plant poisoning investigations.

## 1.1 Introduction

Poisoning by plants causes deaths in livestock every year with accompanying financial losses estimated in the hundreds of millions of dollars (Burrows and Tyrl, 2012; Holechek, 2002). Many losses have been associated with over grazing livestock on rangelands where naturally occurring toxins are found in both native and introduced plant species. Free standing poisonous plants are in general, avoided by livestock in favor of more palatable forages (Stegelmeier and Panter, 2012). Drought and over grazing, however, can result in such high defoliation of preferred species, that photosynthetic capacity is reduced to a point where plants cannot regenerate efficiently and poisonous species either increase in abundance due to niche expansion or simply become more frequently available foraging options. Prepared and stored feeds, fodder, silage, and hay may also be contaminated by toxic plants and fungal metabolites intermingled with crops at time of harvest (Pfister, 1999). Typically, toxic plants are not distributed uniformly across forage crops, so contamination is often patchy, making diagnostics more challenging. Additional identification challenges result from the destruction of characteristic plant structures during feed and fodder production, which hinders morphologically based identification in response to poisoning events (Stegelmeier and Panter, 2012).

Several poisonous plants have played significant roles in livestock deaths from forage consumption in recent years in the Western United States. Some of those include; *Senecio jacobea* (Ragwort) and *Cynoglossum officinal*e (Hounds-tongue) containing pyrrolizidine alkaloids (Baker et al., 1989; Giles, 1983), *Datura stramonium* (Jimsonweed) containing tropane alkaloids (Binev et al., 2006; Naude et al., 2005), *Adonis aestivalis* (Pheasant’s Eye) containing cardenolides(Woods et al., 2004), and *Salvia reflexa* (Lanceleaf sage), containing salviarin. Recently, *Salvia reflexa* was implicated in several catastrophic poisoning events, with one case resulting in 165 cattle deaths (Panter et al., 2021; Stonecipher et al., 2024). In each case, *Salvia reflexa* was a contaminant in newly seeded alfalfa hay (Figure 1).

**Figure 1.**
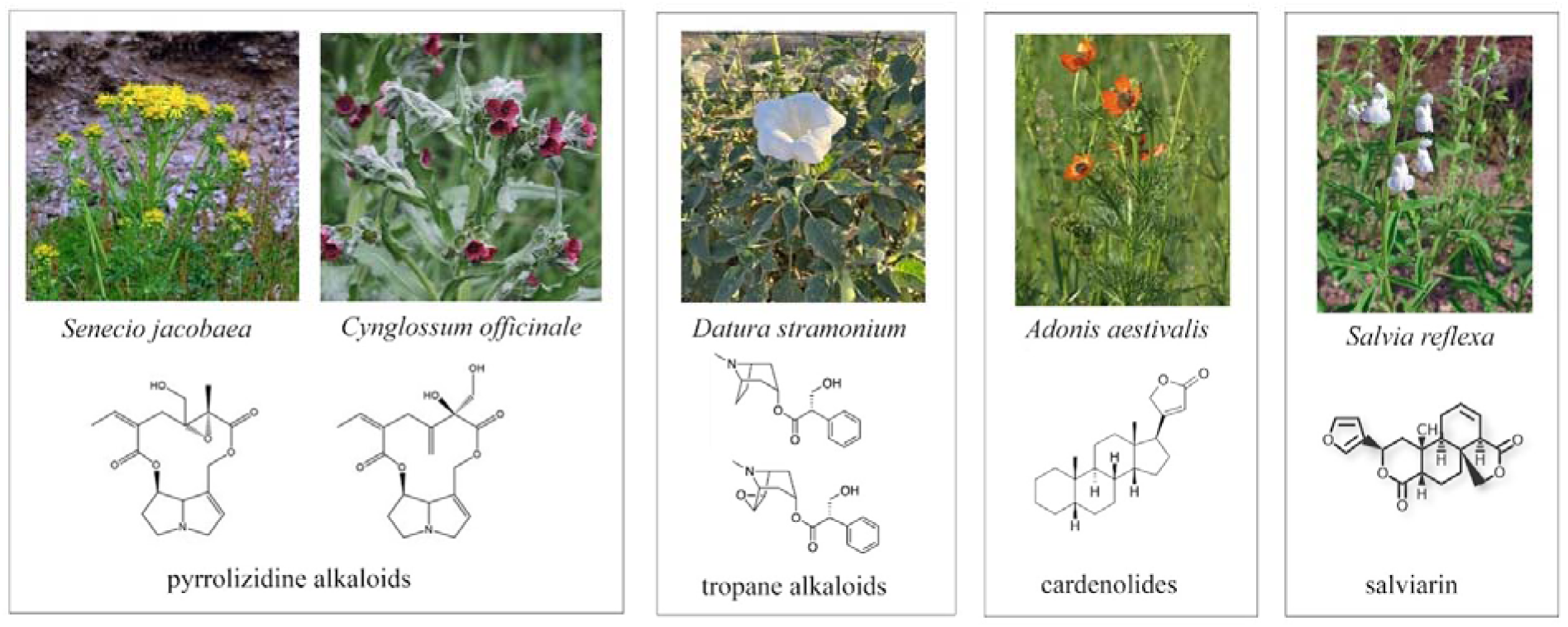
Poisonous plants implicated in recent forage contamination in western states.

Metagenomic data, genome skims and metabarcodes, have demonstrated utility for identification and characterization of plants comprising diet composition of animals (Craine et al., 2016; Craine et al., 2015; Erickson et al., 2017; Scasta et al., 2019; Scasta et al., 2020), herbal products (McShea et al., 2019; Raclariu et al., 2017; Seethapathy et al., 2019) and companion animal kibble (Ottesen et al., 2024). These approaches can both identify plant species and estimate relative abundance of plant components within a given matrix. Erickson et al, for example, used metabarcoding to determine that deer preferentially grazed native plant species in Virginia woodlands, work which demonstrated that genetically based reconstruction of diet can be used to infer behavior and ecology of an herbivore (Erickson et al., 2017). Genome skims have demonstrated utility for identification of single species as well as plant mixtures in dietary supplements (Handy et al., 2021) or probiotics. By targeting high-copy-number elements such as chloroplast DNA, mitochondrial DNA, and ribosomal DNA, shallower sequencing can suffice for taxonomic identification and phylogenetic inference. Likewise, the transfer RNA for leucine (trnL, UAA) gene, located within the chloroplast genome, is widely used in plant identification due to its high degree of conservation and the presence of variable regions that provide species-specific genetic markers. The trnL gene is relatively short, making it easy to amplify even from degraded DNA samples using PCR making it suitable for use with herbarium specimens, environmental samples, and forensic applications where DNA quality may be compromised.

A range of targeted and target independent molecular technologies have been applied to veterinary investigative response efforts as diagnostic tools in cases of poisoning (Jumper, 2024; Lee et al., 2020; Schweikle et al., 2023). Here, we apply metagenomic profiling, genome skims and metabarcoding, and chemistry to retrospectively describe the composition of contaminated alfalfa hay from a case of *Salvia reflexa* poisoning that killed 165 cattle (Panter et al., 2021).

## 2.1 Materials and Methods

### 2.1.1 Chemical analysis

Forage plugs from 220 bales of hay were analyzed chemically for the hepatoxic principle, salviarin, known to be present in *S. reflexa* (Panter et al., 2021). For each chemical assay, a 100-mg aliquot of sample was extracted with 5LmL of dichloromethane for 1 h. Samples were filtered, and 1LmL was transferred to autosampler vials for gas chromatography– mass spectrometry detection (GC-MS; 5977 MSD and 7890B gas chromatograph with a split/splitless injector; Agilent Technologies). Samples (1LµL) were injected in the splitless mode (250°C) onto a DB-5MS capillary (30Lm × 0.25Lmm i.d.) column. The column oven was programmed with the following temperature sequence: 100°C (0–1Lmin); 100–200°C (40°C/min); 200–325°C (10°C/min). Transfer line temperature was 275°C. Mass spectrometry data were collected after ionization at 70eV over a range of *m/z* 50–650.

To estimate the level of contamination by *S. reflexa* in core samples, a set of standards were created by mixing ground *S. reflexa* with ground *Kochia scoparia* at levels of 0.78%, 1.56%, 3.12%, 6.25%, 12.5%, 25%, and 50%, *Salvia* in *Kochia*. Peak area (GC-MS) for salviarin (R_t_ = 15.4Lmin) versus % *Salvia* was then used for calibration.

### 2.1.2 DNA extraction for metagenomes and genome skims

DNA data for metagenomic profiling, genome skims, and metabarcoding were recovered from eighteen plugs selected using a stratified sampling of the full 220 bales of hay with 3 random replicate selections at each of 6 contamination levels (0%, 10%, 20%, 30%, 40%, and 50%). Additionally, rumen contents from three fatally poisoned cows were freeze dried and ground prior to DNA extraction (Qiagen Tissue Lyser II at 30 Hz (1800 oscillations per minute) for 1.3 minutes per sample). Replicates of 100 mg of powder were used for DNA extraction. DNA extraction was performed using the Zymo High Molecular Weight DNA extraction kit according to the manufacturer’s specifications including an extra lysing step using PBS and lysozyme.

Metagenomes were made for each of the three replicates of six contamination levels: 0%, 10%, 20%, 30%, 40%, and 50% by grinding three replicates of 2.5 g portions of each level of contaminated hay, control *Salvia* and rumen. Libraries of DNA were created using the Illumina DNA Library Prep Kit according to the manufacturer’s specified protocols and sequencing was performed on a NextSeq 2000 using a high throughput kit.

### 2.1.3 Metagenomic Informatics

Sequence data were screened for quality metrics using Trimmomatic (Bolger et al., 2014) and analyzed using in house FDA pipelines and databases as previously described(Kocurek et al., 2024). Determination of bacterial and fungal composition from shotgun sequencing was conducted using custom C++ programs developed to compile a *k*-mer signature database containing multiple unique 30 bp sequences per species and then identify each read in the input file using the 30 bp probes. For each bacterial or fungal species or subspecies, each non-duplicated 30-mer from a reference whole genome sequence was placed into a database. Any *k*-mers not found in at least 2/3 of a set of additional genome sequences of the same species were removed and *k*-mers found in genomes of other species were removed. Normalization was performed to correct for bias due to differing number of *k*-mers per database entry and results are tabulated as percent of identified reads (contribution to the microbial population of identified species) for each database entry. Results can be expressed by raw hits or as relative abundance. Both are useful when detecting low abundance organisms in complex metagenomes. Annotation was also evaluated using the Cosmos ID cloud based application (CosmosID Metagenomics Cloud, app.cosmosid.com, CosmosID Inc., www.cosmosid.com) with Fungal Database Version 1.2(Inc., 2022). LEfSe (Linear discriminant analysis effect size) by the Huttenhower biobakery (https://github.com/biobakery/biobakery) (McIver et al., 2017) was calculated in the COSMOSID application. LEfSe (Linear discriminant analysis effect size) is an algorithm used for biological biomarker discovery. Features such as genes, pathways, or taxa are identified for each treatment and the non-parametric factorial Kruskal-Wallis (KW) sum-rank test (Kruskal and Wallis, 1952) is used to identify significant differential abundance of specific features. Linear Discriminant Analysis (LDA) is used to estimate the effect size of each differentially abundant feature and rank the feature accordingly. Sequence data annotations were visualized using graphs created by the R Tidyverse package (https://cran.r-project.org/web/packages/tidyverse/index.html).

### 2.1.4 Mycotoxin pathway identification

Metagenomic reads were mapped with Diamond (v2.0.5)(Buchfink et al., 2021) BLASTX (≥95% identity and those ≥90% read coverage) to a database of genes involved in mycotoxin biosynthetic pathways: aflatoxin, deoxynivalenol, nivalenol, ochratoxin, patulin, sterigmatocystin, T2_toxin, tenuazonic acid. Genes were identified and downloaded from the MetaCyc database (Karp et al., 2019) (https://metacyc.org/). Identity of reads aligning to mycotoxin genes was confirmed by BLASTX aligning them to the NCBI nr database online.

### 2.1.5 DNA metabarcoding

DNA was extracted from rumen contents using the Omega Biotek Mag-Bind® Universal Pathogen Core Kit (4x96 Preps) (Cat. No. / ID: M4030-01) according to the manufacturer’s protocol. PCR primers, trnL c and h were used for PCR amplification. Both forward and reverse primers also contained a 5’ adaptor sequence to allow for subsequent indexing. Amplicons were cleaned and a second round of PCR was performed to complete the sequencing library construct, appending Illumina sequencing adapters and indices. Indexed amplicons from each sample were purified and normalized using mag-bind normalization. Library pools were sent for sequencing on an Illumina MiSeq (San Diego, CA).

### 2.1.6 Metabarcoding Informatics

Raw sequence data were demultiplexed using pheniqs v2.1.0 (Galanti et al., 2021), enforcing strict matching of sample barcode indices (i.e, no errors). For each sample, reads were then clustered using the unoise3 denoising algorithm (Edgar, 2016) as implemented in vsearch, using an alpha value of 5 and discarding unique raw sequences observed less than 8 times. Counts of the resulting exact sequence variants (ESVs) were compiled and a consensus taxonomy was assigned to each ESV using a custom best-hits algorithm and reference database of publicly available GenBank sequences (Benson et al., 2005) and Jonah Ventures voucher sequence records. Consensus taxonomy was generated using either all 100% matching reference sequences or all reference sequences within 1% of the top match. The relative abundance of each ESV for a respective sample was calculated by dividing the number of reads for each ESV by the total number of reads from the respective sample. Sequence variants greater than 2% within a single sample were used for downstream analyses. The complete data set of the DNA metabarcoding results are available upon request.

## 3.1 Results and Discussion

Estimated *S. reflexa* contamination in the hay bales using chemistry ranged from not detected to 86 percent contamination as was shown in Panter et al. (2021). Eighteen plugs were selected for further analysis using a stratified sampling protocol where 3 samples were randomly selected where *Salvia* contamination was estimated using chemistry at not detected, 10%, 20%, 30%, 40%, and 50% with a mean of 25.5 ±17.9%. *Salvia reflexa* contamination was determined in these samples using genome skims and DNA metabarcoding. Genome skims estimated *S. reflexa* contamination from 0.3%-80.4 with a mean of 45.2 ±17.9% (Figure 2, Table 1). In comparison, using trnL for metabarcoding, estimates of *S. reflexa* contamination ranged from not detected to 33.6% with a mean of 15.3 ±10.5% (Figure 2, Table 1). Regression analysis showed a strong association between each of the three respective methods where the coefficients of determination (R^2^) ranged from 0.88 to 0.92 respectively.

**Figure 2.**
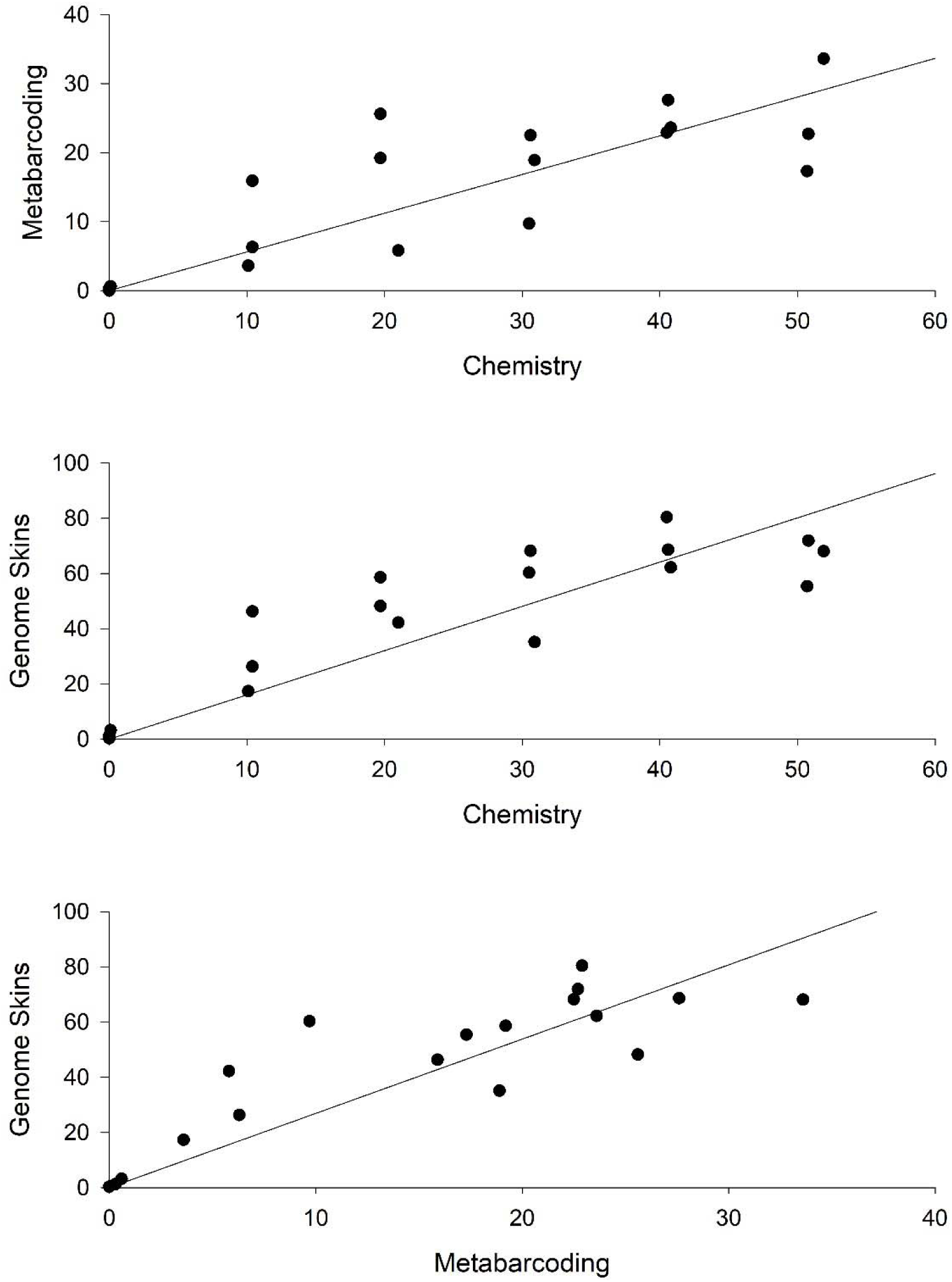
(A) chemistry vs. metabarcoding; GCMS chemical measurement of salviarin plotted against metabarcoding (trnL). (B) chemistry vs. genome skims; GCMS measurement of salviarin plotted against genome skims (k-mer). (C) metabarcoding vs. genome skims; metabarcoding (trnL) vs genome skims (k-mer).

**Table 1.** Range and Means of several plant genera detected among the 18 plugs of contaminated hay. Coefficients of determination (R^2^) comparing estimated composition of a respective plant using genome skims and DNA barcoding.

| | Genome Skims | | DNA Barcoding | | $R^2$ |
| --- | --- | --- | --- | --- | --- |
|  | range (%) | mean (%) | range (%) | mean (%) |  |
| *Salvia* | 0.3-80.4 | 45.2 | nd-33.6 | 15.3 | 0.92 |
| *Amaranthus* | nd-14.5 | 1.6 | nd-6.6 | 0.6 | 0.95 |
| *Bassia* | 1.8-94.0 | 25.9 | 2.5-93.2 | 26.4 | 0.97 |
| *Chenopodium* | 0.1-11.3 | 2.6 | nd-5.6 | 1 | 0.8 |
| *Medicago* | 0.1-45.4 | 14.7 | nd-49.9 | 25.5 | 0.84 |
| *Poaceae* | 0.6-45.3 | 2.5 | 2.1-23.0 | 10.8 | 0.86 |

Visual analysis of the contaminated bales of hay by Panter et al. (2021) tentatively identified the following weedy plants; annual kochia (*Kochia scoparia*; syn. Summer cypress, *Bassia scoparia*), Russian thistle (syn. Russian tumbleweed; *Salsola tragus*), sunflower (*Helianthus* spp.), red root (*Amaranthus* spp.), lamb’s quarter (syn. goosefoot; *Chenopodium* spp.), halogeton (*Halogeton glomeratus*), barnyard grass (*Echinochloa* spp.), cocklebur (*Xanthium strumarium*), pennycress (*Thlaspi arvense*), foxtail (*Setaria* spp.), corn stalks (*Zea mays*), and lance-leaf sage (*S. reflexa)*. No estimates of contamination were performed. Consistent with the visual analysis genome skims and DNA metabarcoding of the representative plugs identified many of the same plant taxa. Among these were an unknown (Asteraceae), *Amaranthus* (Amaranthaceae) *Bassia* (Chenopodiaceae), *Salsola* (Chenopodiaceae), *Chenopodium* (Chenopodiaceae), *Medicago* (Fabaceae), unknown (Poaceae). The relative abundance of each plant genus varied among the different core samples. Relative abundances of the different plant species identified with short read metagenomic data and the k-mer database and pipeline replicates among the 18 different plugs of *Salvia* contamination at six levels: 0%, 10%, 20%, 30%, 40%, and 50% are shown in Figure 3. Similar estimates of the different plant taxa including *Amaranthus, Bassia, Chenopodium, Medicago* and undefined Poaceae (representing several potential taxa) were observed through DNA metabarcoding. Coefficients of determination (R^2^) ranged from 0.80 to 0.97 (Table 1), suggesting a strong association between the two methods in estimating various plant taxa in contaminated mixtures. Notably, genome skims estimated *Salvia*, *Amaranthus, Chenopodium* to have a higher mean amount among the 18 core samples while DNA barcoding estimated *Medicago* and undefined Poaceae to have a higher mean amount among the 18 core samples.

**Figure 3.**
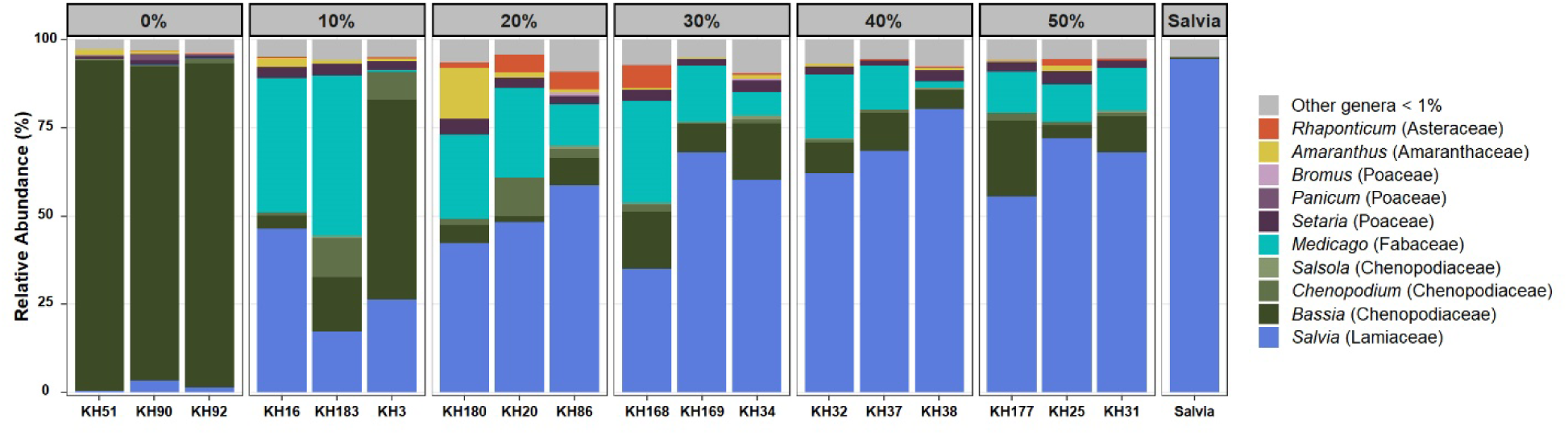
Plant species identified from shotgun DNA of rumen with genome skims (database of unique k-mers from chloroplasts) used for taxonomic ID in 3 replicates of 6 levels (0%, 10%, 20%, 30%, 40%, and 50%) of *Salvia* contaminated hay with one sample of straight *Salvia*.

**Figure 4.**
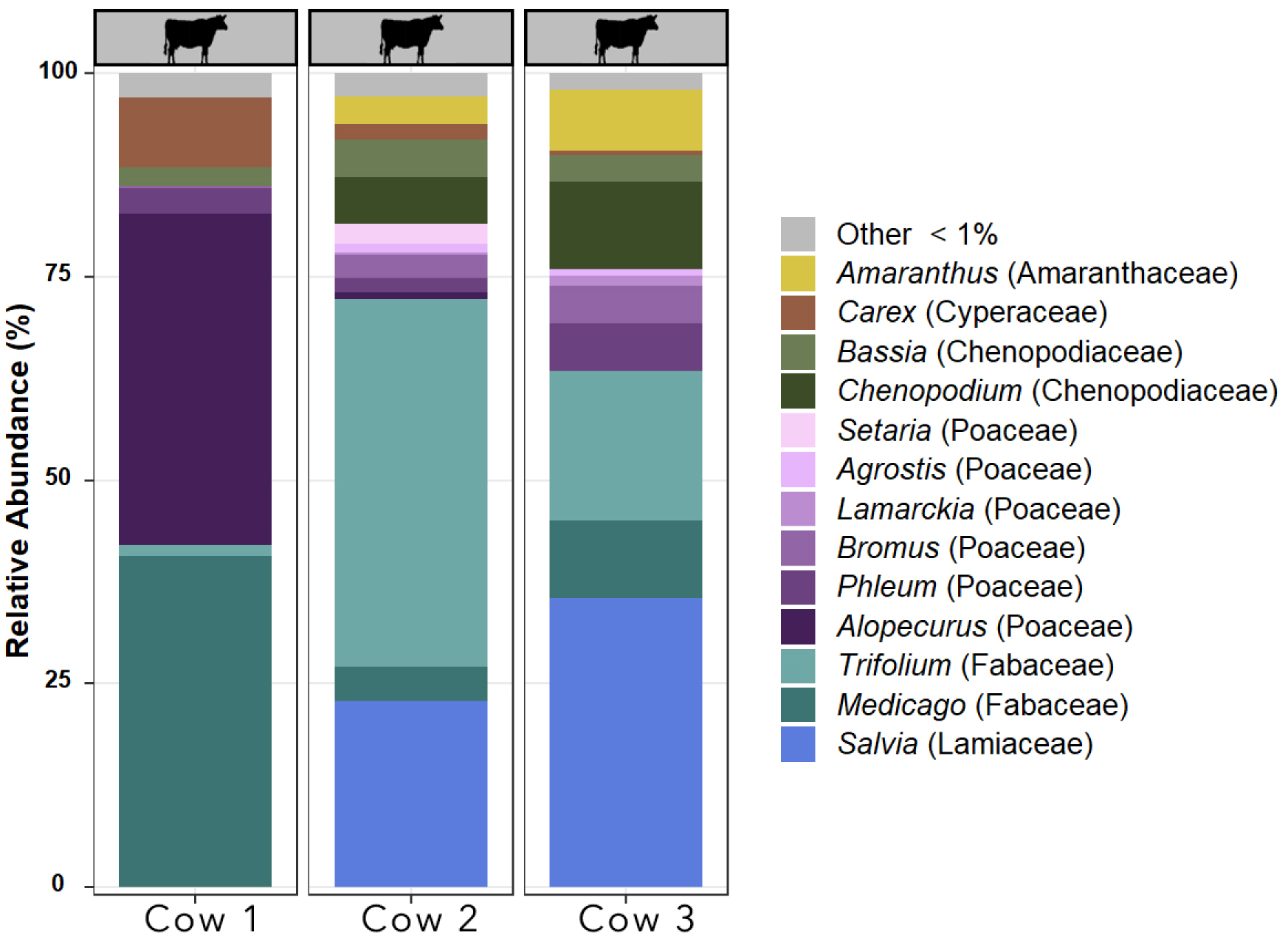
Plant species identified from shotgun DNA of rumen with genome skims (database of unique k-mers from chloroplasts) used for taxonomic ID in the Rumen of 3 deceased cows associated with the *Salvia reflexa* poisoning case.

**Figure 5.**
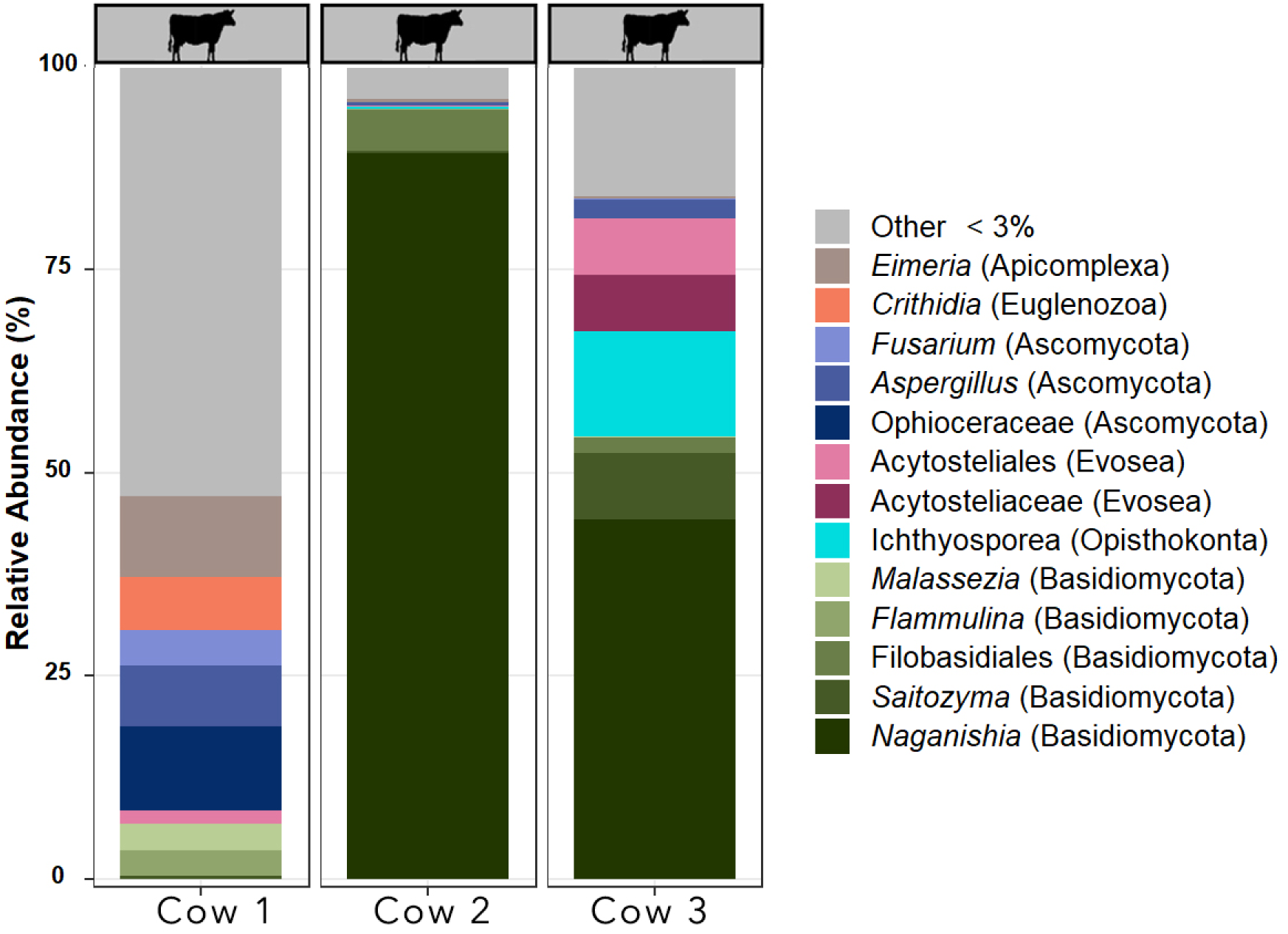
Fungal species identified from shotgun DNA of rumen with metagenomic or genome skims (database of unique k-mers) used for taxonomic ID in the Rumen of 3 deceased cows associated with the *Salvia reflexa* poisoning case.

Rumen contents were available from 3 cows associated with the incidence of poisoning. All three cows were suspected of having died due to *S. reflexa* poisoning. Salviarin was detected by GC-MS in two of the three cow rumen contents. Similarly, *S. reflexa* was detected by genome skimming and DNA barcoding in the same two cows where salviarin was detected but not the third. Relative abundances of the different plant species identified with short read metagenomic data and the k-mer database and pipeline replicates among the three cows is shown in Figure 3. Similar estimated species composition of the rumen contents was observed by genome skims and DNA barcoding. Overall, the genera of plants detected in the rumen contents was similar to the hay composition. Nonetheless, there were other plants detected in the rumen contents that were likely present in the pasture where the cattle were grazing including *Carex*, *Delphinium*, and *Trifolium*. The lack of detection of salviarin and/or *Salvia* in the one deceased cow may be due to variability in the sampling of the rumen contents. The rumen is not a homogenous mixture of the diet of an animal. Alternatively, the *Salvia* consumed may have already moved through the rumen thus making it non-detectable. Likewise, the animal may have ingested another poisonous plant. Larkspur (*Delphinium)*, another known toxic species, was detected in rumen of Cow #1 by genome skims and DNA metabarcoding. Larkspur likely did not contribute to the toxicity of the animals as the animals died in the winter, when larkspur plants are non-toxic. At this point the cattle may have been eating dried stems of various plants in the pasture.

Metagenomic data can describe the total composition (genus, species, subspecies and even serovars and varieties) of plants, animals, insects, bacteria, fungi, viruses, plasmids, and antimicrobial resistance (Sharpton, 2014). Paired chemical and metagenomic shotgun sequencing data is an exciting frontier which will undoubtedly provide valuable information to underpin modernized risk assessment for human and animal foods. For example, fungal profiles for the rumen contents (KHCC and KH811) of the two cows where *Salvia* and salviarin were detected were more similar than KH57 where no salviarin and Salvia were detected. Notably, there were a considerable number of hits to *Aspergillus*, a fungal genus which is known to produce aflatoxin (Klich, 2007) in cow 1 (KH57). Additionally, *Fusarium* was seen in cow 1 which may produce the fumonisins, a mycotoxin (Rheeder et al., 2002). Potentially, these were underlying factors that may have contributed to the death of cow 1.

The significant association between trnL (transfer RNA for leucine) and genome skims take advantage of the multiple copies of plastids in cell thus the significant correlation between the two methods is not surprising. Plant mesophyll cells (the middle layer of a leaf) contain about 20-50 chloroplasts – making this plastid genome highly accessible. The contaminated hay herein, however, has a more complex macrobiome and microbiome – with a large collection of plant, fungal, and bacterial species. Nonetheless, a significant association of the levels of contamination are observed among the 18 core samples from the hay. Furthermore, a significant association was observed between the detection of salviarin by GCMS and the presence of *Salvia*.

In the winter of 2023, there was another documented case of *S. reflexa* poisoning that resulted in the death of 111 cows. In this case we were provided a single forage sample and rumen contents from one deceased animal. Salviarin was detected in the rumen contents and forage (Stonecipher et al., 2024). To explore these methods further, DNA barcoding was used to explore the contents in the forage and rumen contents provided. *Salvia reflexa* was detected in the forage sample and rumen contents by DNA barcoding and salviarin was detected by chemistry (data not shown). This provides further evidence of the utility of these tools to aid in providing a correct diagnosis.

Definitive diagnosis of plant poisoning in grazing livestock is often challenging. Diagnoses are made using a preponderance of many types of data including a history of access to poisonous plants, clinical signs consistent with ingestion of the suspected poisonous plant, histopathology, evidence of plant ingestion by microhistology of rumen contents, detection of the bioactive compounds in gastrointestinal contents and/or animal tissues, or detection of plants in rumen contents by human observation. New methods of metagenomic profiling using genome skims and/or DNA barcoding have utility to enhance diagnostics of plant poisoning cases. Here, we demonstrate the utility of genome skims and/or DNA barcoding to retrospectively investigate the contaminated hay and rumen contents from a case of *S. reflexa* poisoning. The data herein demonstrates that genome skims and DNA barcoding provide another tool to aid in investigating plant poisonings. These technologies provide a global approach to investigate the diet composition of poisoned animals unlike microhistological analysis of plant fragments, which is generally more targeted in terms of plant species (Yagueddu, 1998).

## References

1. Baker, D., Smart, R., Ralphs, M., Molyneux, R., 1989. Hound’s-tongue (Cynoglossum officinale) poisoning in a calf. Journal of the American Veterinary Medical Association 194, 929–930.

2. Benson, D.A., Karsch-Mizrachi, I., Lipman, D.J., Ostell, J., Wheeler, D.L., 2005. GenBank. Nucleic acids research 33, D34–D38.

3. Binev, R., Valchev, I., Nikolov, J., 2006. Clinical and pathological studies of Jimson weed (Datura stramonium) poisoning in horses. Trakia Journal of Sciences 4, 56–63.

4. Bolger, A.M., Lohse, M., Usadel, B., 2014. Trimmomatic: a flexible trimmer for Illumina sequence data. Bioinformatics 30, 2114–2120.

5. Buchfink, B., Reuter, K., Drost, H.-G., 2021. Sensitive protein alignments at tree-of-life scale using DIAMOND. Nature Methods 18, 366–368.

6. Burrows, G.E., Tyrl, R.J., 2012. Toxic plants of north America. John Wiley & Sons.

7. Craine, J.M., Angerer, J.P., Elmore, A., Fierer, N., 2016. Continental-scale patterns reveal potential for warming-induced shifts in cattle diet. PLoS One 11, e0161511.

8. Craine, J.M., Towne, E.G., Miller, M., Fierer, N., 2015. Climatic warming and the future of bison as grazers. Scientific Reports 5, 16738.

9. Edgar, R.C., 2016. UNOISE2: improved error-correction for Illumina 16S and ITS amplicon sequencing. BioRxiv, 081257.

10. Erickson, D.L., Reed, E., Ramachandran, P., Bourg, N.A., McShea, W.J., Ottesen, A., 2017. Reconstructing a herbivore’s diet using a novel rbcL DNA mini-barcode for plants. AoB PLANTS 9.

11. Galanti, L., Shasha, D., Gunsalus, K.C., 2021. Pheniqs 2.0: accurate, high-performance Bayesian decoding and confidence estimation for combinatorial barcode indexing. BMC bioinformatics 22, 1–16.

12. Giles, C., 1983. Outbreak of ragwort (Senecio jacobea) poisoning in horses. Equine veterinary journal 15, 248–250.

13. Handy, S.M., Pawar, R.S., Ottesen, A.R., Ramachandran, P., Sagi, S., Zhang, N.-n., Hsu, E., Erickson, D.L., 2021. HPLC-UV, Metabarcoding and Genome Skims of Botanical Dietary Supplements: A Case Study in Echinacea. Planta Medica 87, 314–324.

14. Holechek, J.L., 2002. Do most livestock losses to poisonous plants result from “poor” range management? Rangeland Ecology & Management/Journal of Range Management Archives 55, 270–276.

15. Inc., C., 2022. CosmosID Metagenomics Cloud. app.cosmosid.com CosmosID Inc.

16. Jumper, W.B., C.; Lee, S.; Cook, D.; Stilwell, J.; Harvey, K.;, 2024. Case Report: Investigating an outbreak of tremorgenic mycotoxicosis in beef cows on pasture in Mississippi due to ergot (Claviceps paspali) infection in Dallisgrass (Paspalum dilatatum). The Bovine Practitioner 58, 59–68.

17. Karp, P.D., Billington, R., Caspi, R., Fulcher, C.A., Latendresse, M., Kothari, A., Keseler, I.M., Krummenacker, M., Midford, P.E., Ong, Q., Ong, W.K., Paley, S.M., Subhraveti, P., 2019. The BioCyc collection of microbial genomes and metabolic pathways.

18. Klich, M.A., 2007. Aspergillus flavus: the major producer of aflatoxin. Mol Plant Pathol 8, 713–722.

19. Kocurek, B., Behling, S., Martin, G., Ramachandran, P., Reed, E., Grim, C., Mammel, M., Zheng, J., Franklin, A., Garland, J., Tadesse, D.A., Sharma, M., Tyson, G.H., Kabera, C., Tate, H., McDermott, P.F., Strain, E., Ottesen, A., 2024. Metagenomic survey of antimicrobial resistance (AMR) in Maryland surface waters differentiated by high and low human impact. Microbiol Resour Announc 13, e0047723.

20. Kruskal, W.H., Wallis, W.A., 1952. Use of Ranks in One-Criterion Variance Analysis. Journal of the American Statistical Association 47, 583–621.

21. Lee, S.T., Welch, K.D., Stonecipher, C.A., Cook, D., Gardner, D.R., Pfister, J.A., 2020. Analysis of rumen contents and ocular fluid for toxic alkaloids from goats and cows dosed larkspur (Delphinium barbeyi), lupine (Lupinus leucophyllus), and death camas (Zigadenus paniculatus). Toxicon 176, 21–29.

22. McIver, L.J., Abu-Ali, G., Franzosa, E.A., Schwager, R., Morgan, X.C., Waldron, L., Segata, N., Huttenhower, C., 2017. bioBakery: a meta’omic analysis environment. Bioinformatics 34, 1235–1237.

23. McShea, W.J., Sukmasuang, R., Erickson, D.L., Herrmann, V., Ngoprasert, D., Bhumpakphan, N., Davies, S.J., 2019. Metabarcoding reveals diet diversity in an ungulate community in Thailand. Biotropica 51, 923–937.

24. Naude, T., Gerber, R., Smith, R., Botha, C., 2005. Datura contamination of hay as the suspected cause of an extensive outbreak of impaction colic in horses: clinical communication. Journal of the South African Veterinary Association 76, 107–112.

25. Ottesen, A., Kocurek, B., Reed, E., Commichaux, S., Mammel, M., Ramachandran, P., McDermott, P., Flannery, B.M., Strain, E., 2024. Paired metagenomic and chemical evaluation of aflatoxin-contaminated dog kibble. Frontiers in Veterinary Science 11.

26. Panter, K.E., Stegelmeier, B.L., Gardner, D.R., Stonecipher, C.A., Lee, S.T., Kitchen, D., Brackett, A., Davis, C., 2021. Clinical, pathologic, and toxicologic characterization of Salvia reflexa (lance-leaf sage) poisoning in cattle fed contaminated hay. Journal of Veterinary Diagnostic Investigation 33, 538–547.

27. Pfister, J.A., 1999. Behavioral strategies for coping with poisonous plants.

28. Raclariu, A.C., Paltinean, R., Vlase, L., Labarre, A., Manzanilla, V., Ichim, M.C., Crisan, G., Brysting, A.K., de Boer, H., 2017. Comparative authentication of Hypericum perforatum herbal products using DNA metabarcoding, TLC and HPLC-MS. Scientific reports 7, 1291.

29. Rheeder, J.P., Marasas, W.F., Vismer, H.F., 2002. Production of fumonisin analogs by Fusarium species. Appl Environ Microbiol 68, 2101–2105.

30. Scasta, J., Jorns, T., Derner, J., Lake, S., Augustine, D., Windh, J., Smith, T., 2019. Validation of DNA metabarcoding of fecal samples using cattle fed known rations. Animal Feed Science and Technology 255, 114219.

31. Scasta, J., Jorns, T., Derner, J.D., Stam, B., McClaren, M., Calkins, C., Stewart, W., 2020. Toxic plants in sheep diets grazing extensive landscapes: Insights from Fecal DNA metabarcoding. Livestock Science 236, 104002.

32. Schweikle, S., Häser, A., Wetters, S., Raisin, M., Greiner, M., Rigbers, K., Fischer, U., Pietsch, K., Suntz, M., Nick, P., 2023. DNA barcoding as new diagnostic tool to lethal plant poisoning in herbivorous mammals. Plos one 18, e0292275.

33. Seethapathy, G.S., Raclariu-Manolica, A.-C., Anmarkrud, J.A., Wangensteen, H., de Boer, H.J., 2019. DNA metabarcoding authentication of ayurvedic herbal products on the European market raises concerns of quality and fidelity. Frontiers in plant science 10, 68.

34. Sharpton, T.J., 2014. An introduction to the analysis of shotgun metagenomic data. Frontiers in Plant Science 5.

35. Stegelmeier, B., Panter, K., 2012. Poisonous Plants and Plant Toxins That Are Likely to Contaminate Hay and Other Prepared Feeds in the Western United States. Rangelands 34, 2–11.

36. Stonecipher, C.A., Gardner, D.R., Webb, B.T., Laegreid, W., Welch, K.D., Stegelmeier, B., gt, Cook, D., 2024. Case Report: Salvia reflexa-contaminated hay poisoning in cattle. The Bovine Practitioner 58, 63–68.

37. Woods, L., Filigenzi, M., Booth, M., Rodger, L., Arnold, J., Puschner, B., 2004. Summer pheasant’s eye (Adonis aestivalis) poisoning in three horses. Veterinary pathology 41, 215–220.

38. Yagueddu, C.C., M.S.; Lopez, T., 1998. Microhistological analysis of sheep gastro-intestinal content to confirm poisonous plant ingestion. Rangeland Ecology & Management/Journal of Range Management Archives 51, 655–660.

